# Cyclic Hypoxia drives Wnt5a-ROR1 stroma-tumor crosstalk promoting breast cancer growth

**DOI:** 10.64898/2026.10.11.758391

**Authors:** Yamin Liu, Shaofei Liu, Sarthak Sarhoo, Xirui Wang, Wenqiang Du, David D Ellison, Junaid Afzal, Yasir Suhail, Mohit Kumar Jolly, Kshitiz

## Abstract

Tumor hypoxia can be stable, or cyclic based on proximity to vasculature, and affects not just the cancer cells, but also other cells in the tumor microenvironment (TME). Although cyclic hypoxia can affect cells in the tumor microenvironment differently from stable hypoxia, its effect on non-malignant cells in the tumor microenvironment (TME) is unclear. Here, using single cell RNA ATAC sequencing of multicellular co-culture model of triple-negative breast cancer (TNBC), comprising MDA-MB-231 cells, fibroblasts (Fbs), mesenchymal stromal cells (MSCs), endothelial cells, and macrophages, we systemically map the transcriptomic response shaped by dynamic oxygen regimes. Cyclic hypoxia induced a distinct transcriptional program compared to stable hypoxia in a cell type specific manner, regulating cell subpopulation proportions with key phenotypic implications. In TNBC, cyclic hypoxia reversed chronic hypoxia-induced cell cycle arrest, and reversed the enrichment of a highly mesenchymal WNT5A+/COL6+ subpopulation stabilized in hypoxia. Furthermore, cyclic hypoxia increased fibroblast activation, but suppressed MSC proliferation. Mechanistically, cyclic hypoxia coordinated paracrine WNT5A/B-ROR1 signaling between fibroblasts and TNBCs. In a mouse breast CAF-TNBC co-xenograft model, disrupting this cyclic hypoxia driven WNT5A-ROR1 stroma-cancer crosstalk reduced tumor growth *in vivo*. Together, our findings highlight dynamics in oxygen regimen as distinct microenvironmental cues, reprograming the TME compartments to accelerate cancer growth and invasion.

## Introduction

Hypoxic regions develop in solid tumors due to an imbalance between oxygen supply and consumption during rapid cell growth^1,2^. Low oxygen tension stabilizes Hypoxia Inducible Factor-1 (HIF-1), a heterodimeric complex consisting of molecular oxygen regulated HIF-1α subunit, and a ubiquitous HIF-1β subunit. HIF-1 orchestrates cellular adaption to hypoxic stress primarily by transcriptional activation of a large number of genes^3–5^, as well as by non-transcriptional roles including induction of cell cycle arrest^6,7^. Tumor hypoxia is often spatially, and temporally heterogeneous^8^. Peripheral hypoxic regions in the cancer microenvironment can experience unstable, or oscillatory/cyclic hypoxia owing to leaky neovasculature, or lack of temporal homeostasis in oxygen demand and supply in the rapidly proliferating tumor^9–11^. In addition to instability in hypoxia itself, HIF-1 may also exhibit cyclic activity due to degradation induced by accumulated lactate, itself a product of HIF-1 driven glycolysis^12,13^.

In breast cancers, hypoxia is a common and consequential environmental factor^14,15^, and bulk analysis of breast cancer cells have shown that oscillatory, or cyclic hypoxia can permit escape from HIF-1 induced arrest on cellular growth^12^, and increased expression of pro-tumorigenic and anti-survival genes^16–18^. Although, both sustained and cyclic hypoxia affect the whole TME, considerably little is known of the effect of cyclic hypoxia on the non-malignant cellular component of TME.

To address this question, we established a multicellular co-culture model of triple-negative breast cancer (TNBC) microenvironment in dynamically perfused oxygen tension. The model incorporates MDA-MB-231, a triple negative breast cancer cell line, BJ fibroblasts (Fbs), HUVECs, an endothelial cell line, bone marrow derived mesenchymal stromal cells (MSCs), which are known to be recruited to breast TME and likely contribute to CAF pool^15^, as well as macrophages (Mϕ) differentiated from peripheral blood derived monocytes. Using integrated single cell RNA and ATAC sequencing, we systematically mapped cell-type specific transcriptional responses in stable, and cyclic hypoxia conditions. Stable hypoxia resulted in distinct cellular responses across cell populations, characterized by increased glycolysis, inhibition of cell cycle in most cells but an increase in MSCs, and downregulation of inflammatory polarization in macrophages. In contrast, cycling hypoxia partially reversed several chronic hypoxia driven changes, while inducing unique transcriptional states not observed under chronic hypoxia. Importantly, these specific changes in cyclic hypoxia were driven by emergence of cellular subpopulations in cyclic hypoxia with distinct gene expression signature. These changes included increased mesenchymal transition in malignant cells, enhanced fibroblast activation response in Fbs and MSCs, and increased M2-like polarization in macrophages. Furthermore, cyclic hypoxia selectively remodeled paracrine communication between breast CAFs and MDA-MB-231 via WNT5A and WNT5B signaling via ROR1, enhancing cancer growth and dissemination potential. Overall, our findings reveal that cyclic hypoxia is not merely an intermediate form of chronic hypoxia, but a distinct microenvironmental state that actively rewires stroma-tumor interactions enhancing pro-tumorigenic signaling beyond the effect of chronic/stable hypoxia.

## Methods

### Cell lines

MDA-MB-231 (ATCC #HTB-26) cells were purchased from American Type Culture Collection (ATCC) and cultured in Leibovitz’s L-15 Medium (ATCC 30-2008) supplemented with 10% fetal bovine serum (Gibco), and 1% Pen/Strep in an incubator at 37 °C and 5% CO_2_. Human Umbilical Vein Endothelial Cells (HUVECs) were purchased from LONZA (Cat# CC-2935) and cultured using VascuLife® VEGF Endothelial Medium Complete Kit (Cat# LL-0003, Lifeline cell technology) with 1% Pen/Strep. BJ cells were purchased from ATCC. Bone marrow derived mesenchymal stromal cells (BM-MSCs), is kindly provided by Dr. Kumbar laboratory. Cell cultured in Mesenchymal Stem Cell Growth Kit (ATCC: Cat # PCS-500-040). Human monocytes are isolated from peripheral blood monocyte cells (PBMCs). And differentiated into unpolarized macrophages (MФ differentiated to M0 phase). Cancer-associated fibroblasts (CAFs) isolated from breast cancer biopsy (bCAF), were maintained in DMEM/F12 with 10% FBS, 1% 1% Pen/Strep.

### Monocytes isolation and differentiation

Whole blood from anonymous health donors were obtained from UConn Health Biospecimen Core. The PBMCs were isolated from human whole blood using density gradient centrifuge tube (SepMate™-15, stemcell) following the manufacturer’s protocol. Briefly, 40 mL of fresh human whole blood in Acid-citrate-dextrose (ACD) anticoagulant vacuum tubes were diluted with equal volume of phosphate-buffered saline (PBS) with 2% fetal bovine serum (FBS). The mixture was then transferred to SepMate tube. Centrifugation was performed at 1200 x *g* for 10 minutes at room temperature. After isolation, cells were washed twice in PBS with 2% FBS.

Subsequently, Cells will be isolated from PBMCs using immunoselection isolation monocytes kit (Milenyi Biotec) according to the manufacturing protocol. Briefly, PBMCs were diluted to 107 cells in 30 μl buffer and incubated with 10 μl Biotin-antibody cocktail with 10ul of FcR Blockig Reagent for 5 min in the refrigerator. Following the addition of 20 μl/ml Anti-Biotin MicroBeads, the cells are incubated for 10 min at refrigerator. PBMCs were then placed in the ‘MidiMACS’ magnet Separator (Miltenyi Biotec) for 3 min, The supernatant containing monocyte cells were collected and wash in PBS (with 2% FBS) to obtain pure monocytes using published protocol.

Purified monocytes will be cultured in the RPMI (ThermoFisher Scientific) with 10% heat-inactivated FBS (Hi-FBS), 1% penicillin/streptomycin (ThermoFisher Scientific), with 10 ng/ml Macrophage colony-stimulating factor (M-CSF) (Peprotech) and 1 ng/ml GM-CSF (Peprotech) for 5 days for to differentiate into monocyte-derived macrophages (MDMs) ^19^. Media were changed every 2-3 days.

Our data demonstrate the effectiveness of this method in obtaining pure M0 MACs as the cells exhibit the characteristic morphology with ∼99% cells express CD14 and CD16 markers and only 3% of the cells show M1 marker CD80 and 0.5% M2 marker CD206; Furthermore, after stimulation with 100 ng/mL lipopolysaccharide (LPS), and 20 ng/mL interferon gamma (IFN-γ) stimulations for 12h, the MACs exhibit the characteristic morphology of M1 macrophages.

### Multiome Library Preparation and Sequencing

All the cells, except the macrophages were passaged before reaching 70% confluency at 1:10 ratio. The cells were dissociated and co-cultured on a gas permeable plate (Coy Laboratory Products) at the following ration: MDA-MB-231:Fb:HUVEC:MSC:M0 = 40:40:10:5:5. The co-cultured cells were maintained in a treatment medium, which contains no glucose DMEM (Gibco, ca#A1443001), MEM Non-essential Amino Acid (NEAA) (Cat# 11140-050, Gibco, 1:100), sodium pyruvate (1mM) (Cat# 11360-070, Gibco, 1:100), HEPES Buffer (Ca# BP-299, ThermoFisher,1:100), 10 mM glucose; Insulin-Transferrin-selenium (ITS-G) (#41400045,1%, ThermoFisher); 2% Fetal Bovine Serum (FBS) ; 2% Bovine serum albumin (BSA), 1%pen/strep. The mixed cells were first co-cultured in the treatment medium for 24 h, then subjected to in a normal, hypoxic, or cyclic hypoxia atmosphere for 48 h. For control normoxia condition, cells were cultured in a normal atmosphere. Under hypoxia, cells were incubated in a hypoxia incubator chamber (Embrient Inc). Under intermittent hypoxia, cells were subjected to repeated hypoxia (1 h, 1% O2)/reoxygenation (30 min, 20% O_2_) cycles with in a O_2_ Control InVitro Cabinet (Coy Laboratory Products).

Nuclei were extracted from cell suspensions using 6 minutes lysis time for all samples and otherwise following 10x Genomics Demonstrated Protocol CG000365, Rev C. Isolated nuclei were washed and resuspended in nuclease-free water + 1mM DTT + 25U RNase inhibitor (Sigma Protector, 3335402001). Nuclei quality was assessed via brightfield imaging and Trypan Blue staining, and nuclei concentration was measured by AOPI staining using a Luna FX7 (Logos Biosystems). Following User Guide CG000338 Rev F, chromatin was transposed for 60min at 37C and immediately processed as follows. Approximately 12,000 nuclei from each sample were loaded onto one lane of a 10x Genomics Chip J, and single nucleus encapsulation, barcoding, and library preparation were performed using the Chromium X platform (10x Genomics) according to the manufacturer’s user guide. Resulting libraries were assess for quality via Agilent Tapestation 4200, and quantified by KAPA qPCR and Qubit fluorometry. Libraries derived from mRNA and chromatin were pooled separately and sequenced on separate lanes of a NovaSeq X Plus 10B 200 cycle flow cell (Illumina) targeting 420M read pairs per library using a 50-10-24-90 asymmetric paired-end read configuration (R1-I1-I2-R2).

Illumina base call files for all libraries were demultiplexed and converted to FASTQ files using Illumina bcl2fastq 2.20.0.422. Digital cell-by-gene and -fragment count matrices were generated for each paired set of mRNA- and chromatin-derived libraries against the 10x Genomics GRCh38 reference assembly (version 2020-A; assembly: Ensembl release 98 ; annotations: Gencode v32) using 10x Genomics cellranger-arc count version 2.0.1 pipeline.

### Bioinformatics

The RNA part of the single cell multiomic data was normalized by scTransformwith dimensionality reduction performed using Principal Components Analysis (PCA), and 50 PCA dimensions were used to visualize the data with UMAP in Seurat 5.0.1^59^. ATAC data was processed with Signac 1.12.0^60^, and also visualized in UMAP using 50 PCA dimensions. The RNAseq and ATACseq components of the data were merged in a weighted form by WNN^61^ to do be visualized in UMAP. Canonical markers were used to identify fibroblast, malignant epithelial (tumor), mesenchymal stem cell (MSC), macrophage and endothelial cell populations based on RNA level data. The cells were also tracked for the Control, Lactate, Hypoxia and Oscillation conditions from which they were sequenced. Top 10 genes for each cell type are plotted as heatmap in Fig. 1H for illustrative purposes. For each cell type, subclusters were obtained using the Louvain algorithm with resolution 0.5 for MSCs (4 subclusters), malignant_epithelial (13 subclusters), macrophage (4 subclusters), fibroblast (13 subclusters), and endothelial (2 subclusters). For each celltype, condition wised heatmap were based on significant top 3 hallmark pathway (the two comparisons H vs N and O vs N), and top 3 leading edge genes were chosen based on the log fold change, and then merged for each comparison for visualization. EMT and pEMT scores were computed at the single-cell level using the AUCell algorithm on normalized gene expression^20^. The Hallmark EMT gene set from MSigDB was used to quantify EMT activity^21^, while the pEMT gene signature described by Puram et al. was used to assess partial EMT^22^.

**Figure 1.**
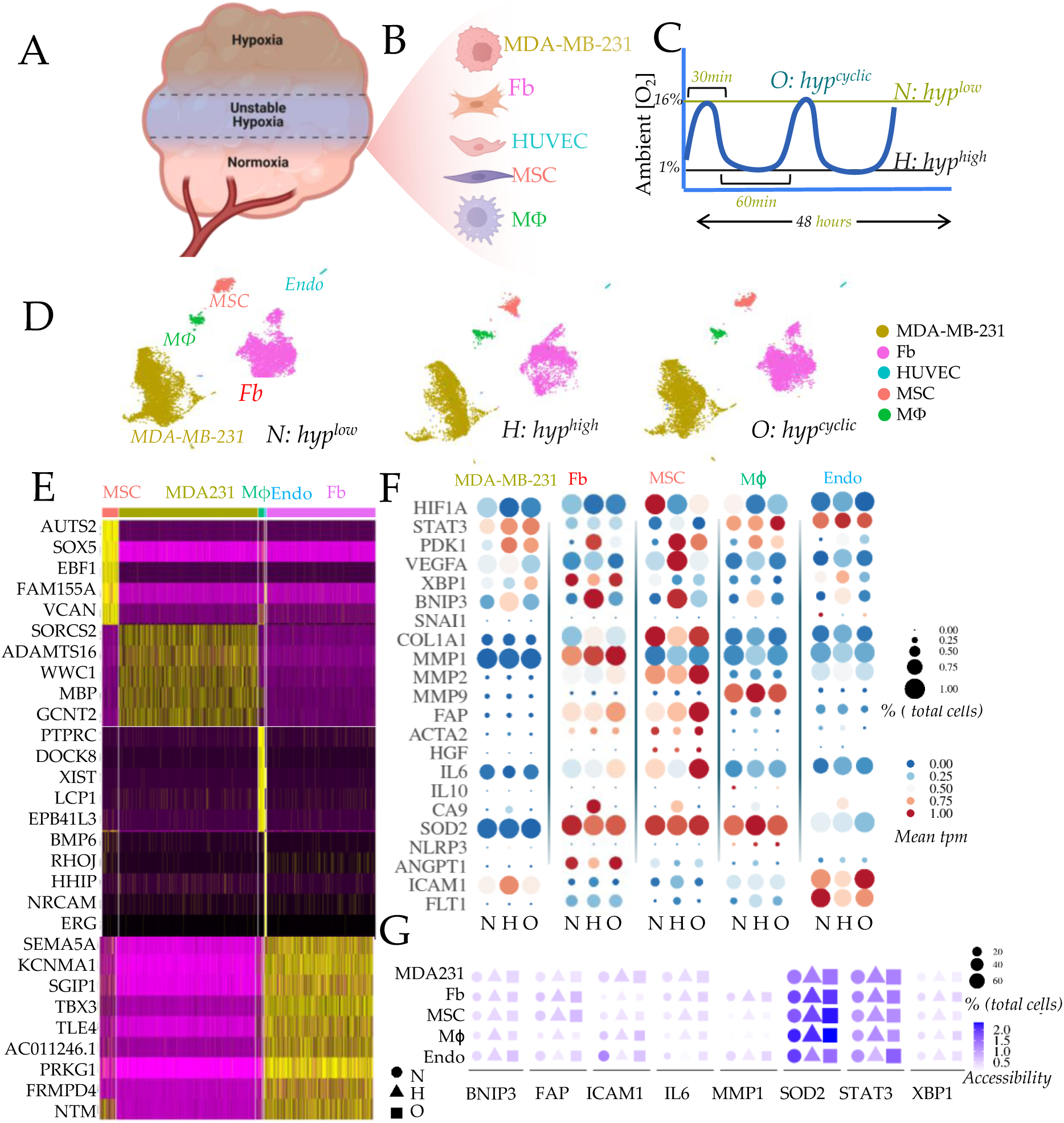
Single cell multiomic analysis of cells in TNBC-TME in stable and cyclic hypoxia. (A) Zones of oxygen deprivation in poorly vascularized tumors can lead to regions of unstable hypoxia distal to vasculature; (B-C) Schematic showing experimental plan, with 5 different tumor associated cells: MDA-MB-231 TNBC cell line, Fibroblasts (Fb), Mesenchymal stromal cells (MSC), non-polarized macrophages differentiated from PBMCs (Mϕ), and endothelial cells (HUVEC), co-cultured *in vitro* on an oxygen permeable plate (C) treated with directly bioavailable stable, or cyclic hypoxia, and normoxia. (E) t-SNE plots for co-cultured cells, identified in separated clusters; (F) Heatmap showing relative expression of top differential genes identified for each cell type; (G) Bubbleplot showing expression of key genes critical for the function of the co-cultured cell types showing differential expression in stable, or fluctuating hypoxia; size: percentage of expressing cells, color denote mean relative expression; (H) Chromosomal accessibility of cis-regulatory region for selected genes in each cell type in N, H, O.

### RNA isolation and cDNA synthesis

Total RNA from MDA231 or Cancer fibroblasts was extracted using an RNA isolation Kit (Life Technologies) according to the manufacturer’s instructions. The amount of each samples’ total RNA was detrimental using a NanoDrop 1000 spectrophotomereter (Thermo Scientific). Total RNA (1 μg of each sample) was reverse-transcribed into cDNA using cDNA Reverse Transcription kit (Applied Biosystems).

### qRT-PCR

Real-time quantitative reverse transcription polymerase chain reaction (qRT-PCR) for mRNA was run on a Bio-Rad CFX Real-Time qPCR Thermal cycler (Bio-rad). Quantitative PCR was carried out at 95°C for an initial 3 min followed by 40 cycles of denaturation at 95°C for 15 s, annealing at 60°C for 30 s and extension at 72°C for 30 s using SYBR^®^ Green PCR Master (Applied Biosystems, CA, USA). Each assay was performed in duplicate, and glyceraldehyde-3-phosphate dehydrogenase (GAPDH) was used as the internal control. Relative gene expression levels were quantified with 2^-ΔΔCT^ -method compared to GAPDH.^62^

The sequence of following primers were used; The qRT-PCR primers are listed in Table 1.

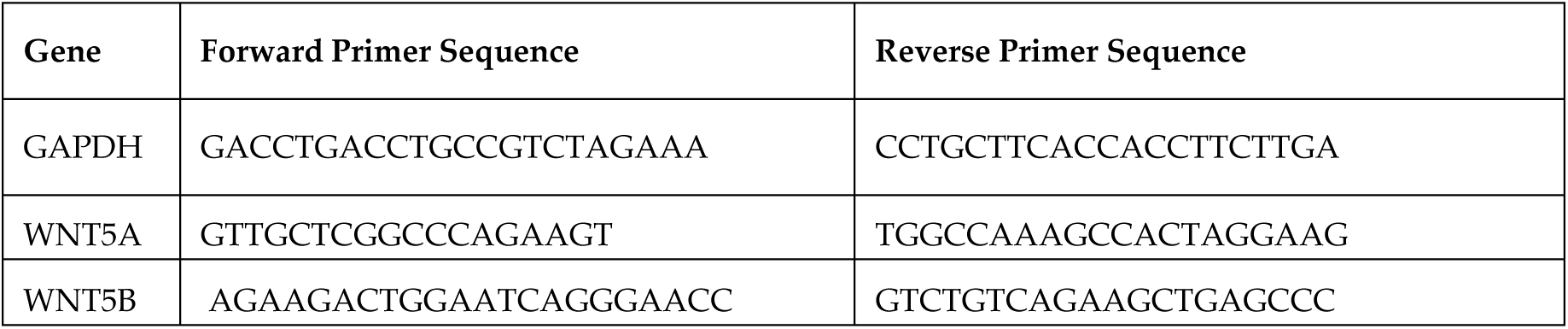
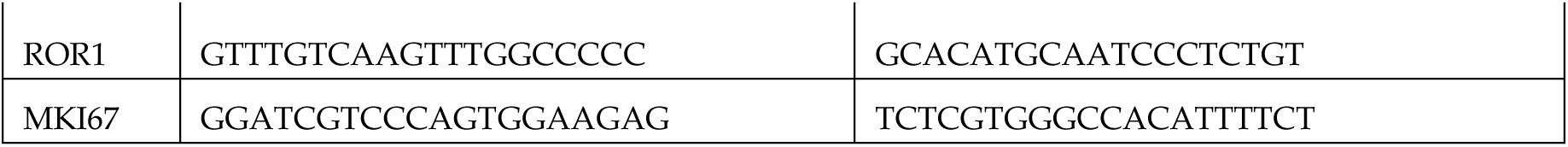

### Virus transduction and selection of cell line

MDA-MB-231 cells at 50-75% confluent, were transduced with Hsp70-Firefly Luc (Puro) Lentivirus (cat# LVP570-PBS, GenTarget Inc) expressing both firefly luciferase and GFP. After 2d, transduced cells were then selected with puromycin and surviving colonies were screened for bioluminescence in complete media supplemented with 150 μg/ml D-luciferin (Cat# LUCCK-100; Goldbio) by *in vitro* imaging using the IVIS camera system (Ami HT Spectrum, Arizona, USA). Bioluminescent, antibiotic resistant, and GFP cell clones were amplified in culture and characterized for stable luminescence *in vitro* and tumorigenic potential *in vivo*.

### Gene silencing

ROR1 silencing in MDA-MB-231 cells for *in vitro* experiments: MDA-MB-231 cells were transfected with human DsiRNA ROR1 and Negative control using Lipofectamine RNAiMAX reagent (LifeTechnologies) according to the manufacturer’s protocol for 48 h, then the transfected cells are used for next step assay. Their sequences were as follows: Negative control siRNA (# 51011403, IDT); ROR1 siRNA, 5’-GUCACUUAUCUGAUAAGU -3’ (sense) and 5’-UGAGAACACUUAUCAGAU -3’ (antisense).

Generation of stable ROR1-knockdown cells for *in vivo* experiments: A stable ROR1 knock down MDA-MB-231-Luci-GFP cell line were generated using CRISPR-Cas9 genome editing system and subsequently used for *in vivo* experient. Pre-designed synthetic sgRNAs (IDT, design IDs:# Hs.CAS9.ROR1.1AC; IDs:# Hs.CAS9.ROR1.1AD) and Cas9 protein were transfected into cells using the Neon™ Transfection System (Invitrogen) according to the manufacturer’s instructions. Briefly, on the day of transfection, cells were cultured to 50–70 % confluence. Cells were detached and resuspended at 2 x10^5^ cells in 5 uL Buffer R. CRIPSR-Cas9/gRNA ribonucleoprotein (RNP) complex were prepared by mixing 1 μg of TrueCutTM Cas9 Protein v2 (Invitrogen, cat#36498) and 250 ng of gRNA in 5 μL of Resuspension Buffer R, followed by incubation at room temperature for 10–20 min to allow RNP complex formation. After incubation, the Cas9/gRNA RNP complexes were gently mixed with cell suspension, and the mixture was loaded into the NeonTM 10-μL tip. The electroporation program was performed using program (1600 V/10 ms/3 pulses). Immediately transfer the electroporated cells into a 24-well plate containing 0.5 mL of pre-warmed culture medium. At 48h post-transfection, the cells were harvested for analysis of genome modification efficiency using RT-PCR. The bioluminescence imaging (BLI) intensity of the control and ROR1-knockdown MDA-MB-231-Luci-GFP cell lines was evaluated prior to implantation into mice and showed no significant difference between the two groups.

### Immunostaining

Cultured cells were washed twice with PBS before fixing them with 4% paraformaldehyde (pH 7.4) for 15 min at room temperature. After fixation, cells were washed with cold PBS before permeabilizing them with PBS buffer containing 0.2% Triton X-100 and 0.1% BSA for 15 min followed by one-hour incubation with blocking buffer (5% goat serum in PBS). Cells were then incubated overnight with primary antibodies with subsequent 30 min incubation with secondary antibody. Following primary antibodies were used for labeling; with Ki-67 (Cat.# 27309-1-AP; 1:500; Proteintech), Following Alexa Fluor Secondary Antibodies (ThermoFisher) were used: Alexa Fluor 488 (A32731), Phalloidin (Cat# A12381, Invitrogen) was used to label the cell cytoskeletal protein F-actin. Cells were counterstained with DAPI (ThermoFisher D21490) before imaging. The data was analyzed using ImageJ software (version 1.48, National Institutes of Health, Bethesda, MD, USA).

### Accelerated nanopatterned stromal invasion assay

MDA231 were initially transfected with siCTRL or siROR1 for 24h. After transfection, the cells were labelled with CellTracker™CM-Dil Dye (Thermo Fisher Scientific). The labeled cells were then immediately seeded at a density of 5 X 10^4^ cells per well into a 96 - well NanoSurface Plate (Curibio) and incubated in serum-free medium for another 12h.

To create the wound, an auto scratch machine (Agilent BioTek AutoScratch) is used to create the gap in the 96 well. The wells were then washed twice with PBS to remove the unattached cell. The unlabeled stromal cells (bCAFs) were seeded at a density of 5 x10 to fill and attach the scratched gap. After incubation for 5h, the unattached stromal cells were removed by washing off. Then the medium was replaced with fresh serum-free medium, with or without Wnt5a Wnt5b (#OPCA04542; # OPCD07977, Aviva Systems Biology Corporation). Then the plate is ready to be mounted on the stage of the live-cell microscopy.

MDA231 invasion to stromal cells was performed using a 20X objective on Zeiss Observer Z1 microscope. Cells were images using at 2 h intervals for a total duration of 24h. The underlying anisotropic nanotextured substrate induced highly directional migration protrusions, amplifying the migration process while enabling efficient data analysis due to the reduced dimensionality. The invasion area were tracked and quantified using the Region of Interest (ROI) panel in Fiji software. The ROI selection involved manually tracing the migration boundaries. The extent of the migration area was calculated by 24h ROI area subtracting the 0h ROI area. Each experiment was conducted in 8 replicates. The data were expressed as Mean ± SEM.

### Cell proliferation assay

Cell proliferation was measured using the Cell Counting Kit-8 (Dojindo) or PrestoBlue^TM^ Cell Viability reagent (Thermofisher, USA) according to manufacture’s protocol. Briefly, cells was seeded into the wells of a 96-well microtiter plate (5 X 10^4^ per well) and incubated at 37 °C for 24 h, following for starving 12h, then cells were subjected to different treatment condition at 37 °C for 24 h; CCK-8 solution (10 μL per well) was added, followed by an additional incubation for 4 h. The absorbance was measured at 450 nm. In PrestoBlue^TM^ assay, 10% PrestoBlue^TM^ reagent were added to 96 well with 1h incubation, then measured on the Perkin-Elmer EnVision at 530/595 nm.

### Enzyme-Linked Immunosorbent Assay

MDA231 were seeded on a gas permeable plate (Coy Laboratory Products) in the treatment medium for 24 h, then subjected to in a normal, hypoxic, or cyclic hypoxia atmosphere for 24 h. Supernatants were collected and used for Enzyme-Linked Immunosorbent Assay (ELISA) analysis for quantification of secreted WNT5A and WNT5B (CUSABIO) following the manufacturer’s instructions.

### Mouse strain and animal care

Mice were housed in ventilated cages within an accredited facility under a 12-hour light/dark cycle with temperature-controlled (22 ± 2 °C) and humidity-controlled (55 ± 5%) environment and fed a standard chow diet. All animal protocols were approved by the Institutional Animal Care and Use Committee (IACUC) at the University of Connecticut Health Center before study initiation, and all experiments were performed in accordance with IACUC guidelines. Mice used in these studies were female NBSGW mice (Strain #:026622, JAX lab, USA). Animal experiments were under IACUC compliance: Protocol Number AP-200781 approved in 2024. Sex: Females mice were used for TNBC model for breast fat xenografts. Age: 12-24 weeks. Power calculation was used to determine mouse numbers based on in vitro experimentation.

### Breast cancer mouse model

Breast cancer mouse model was established accordingly to a previously published protocol^23^. Briefly, Female NBSGW mice were anesthetized by exposure to 1–3% isoflurane and orthotopically injected into the mammary fat pad with a Matrigel mixture, which contain 0.15× 106 MDA231-luc-GFP cells with or without ROR1 Knockdown, with 0.15× 106 human cancer associated fibroblasts (CAF), suspended in 100 μl 50% DPBS/50% Matrigel. Tumor growth and metastasis was monitored weekly by the bioluminescent imaging and external caliper measurements (tumor size = [length × width × height] × 0.52) each week.

### Bioluminescent imaging

Bioluminescent imaging was performed with Ami HT Spectrum system (Spectral instruments imaging, Arizona, USA) or Xenogen IVIS 200 Spectrum imaging system (PerkinElmer, USA). For *in vivo* imaging, mice were injected D-luciferin by intraperitoneal at 150 mg/kg in DPBS, and anesthetized (1-3% isoflurane). Mice were then placed onto the warmed stage inside the light-tight camera box with continuous exposure to 1-2% isoflurane. Imagines were taken after 10-15 min post-injection to capture the bioluminescent signals. Imaging times ranged from 1 to 60 s depending on the time point. Generally, 5 mice were imaged at a time. The bioluminescent signals were subsequently visualized and quantified using Aura *in vivo* imaging software (Spectral instruments imaging, Arizona, USA). Background bioluminescence *in vivo* was in the range of 1-2 × 10^5^ photons/s. At the end of the experiments, primary tumor were excised and weighed.

### Statistical Analysis

Statistical analyses were performed with GraphPad Prism version for windows (GraphPad Software, San Diego, CA). Group analyses were conducted with one-way Anova and Tukey’s test. P <0.05 was considered statistically significant.

## Results

### Single cell analysis of transcriptomic changes in breast TME in stable and cycling hypoxia

To model the cellular complexity of breast cancer TME in normoxic, and hypoxic regions – stable, or cyclic (**Figure 1A**), we co-cultured key cell types in triple negative breast cancer TME, sharing the same medium. While Fbs, endothelial cells, as well as tissue resident MФ are part of TME, MSCs as well as MФs are also recruited to breast cancer from bone marrow or peripherally^24,25^, likely to serve their purpose of tissue repair, but also aided by specific HIF induced recruitment signals^15^ (**Figure 1B**). Oxygen has very low permeability in aqueous solutions^26^, therefore temporal dynamics in atmospheric oxygen level is not rapidly equilibrated within the culture medium. We therefore co-cultured cells on a gas permeable plate controlling ambient oxygen tension with a dynamic oxygen perfusion system. Ambient oxygen is therefore directly sensed by the adherent cells through the bottom of the plate.

We performed single cell RNAseq and ATACseq (multiomic) analysis after 48 hours of treatment with normoxia, hypoxia, or a regimen of cycling hypoxia matching observed oxygen transients in animal models (**Figure 1C**)^27,28^. Successfully sequenced cell counts for each cell type are listed in **Figure S1A**. Cell clusters were well separated based on both ATACseq and scRNAseq (**Figure 1D**) with distinct gene expression patterns (**Figure 1E**). We computed the relative expression of genes critical to above cell types in hypoxia in the three regimens: normoxia (N), stable/sustained hypoxia (H), cyclic/oscillatory hypoxia (O) (**Figure S1, S2**). For most genes, cycling hypoxia expectedly resulted in a sub-hypoxic response^29^, in between normoxia and hypoxia. However, exceptions were notable in some of the key genes where cycling hypoxia resulted in a specific pattern of expression (**Figure 1F**). Among the most notable cyclic hypoxia-specific changes were increased expression of STAT3 in MФ, a key regulator of pro-tumorigenic M2 polarization^30^, and elevated expression of XBP1 in MDA-MB-231, a central mediator of the unfolded protein response (UPR). In the stromal populations, both Fbs and MSCs showed key marker of fibroblast activation elevated in cyclic hypoxia. MMP1, IL6, as well as fibroblast activation protein (FAP) were increased and BNIP3 decreased in both stromal cell types, while MMP2 increased in MSCs. Endothelial cells similarly showed increased expression of ICAM1, which encodes intercellular adhesion molecule-1. Overall, the biological effect of cyclic hypoxia appears to be elevating the pro-tumor cellular microenvironment, underscored by the established role of XBP1 mediated UPR in drug resistance^31^, matrix metalloproteinases in extracellular matrix remodeling and invasion^32^, ICAM1 in leukocyte adhesion for extravasation^33^, and stromal FAP expression associated with poor clinical outcomes^13^. ATACseq for selected genes for each cell type also highlighted that cyclic hypoxia has a distinct effect on transcriptional regulation via regulation of chromosomal accessibility for many genes (**Figure 1G**). These observations prompted a systemic investigation of the cell-type specific transcription uniquely induced by stable vs cyclic hypoxia.

### Cycling hypoxia rescues proliferation from hypoxic growth arrest, and reverses enrichment of WNT5A/COL6+ subpopulation enrichment

We first examined the effects of chronic and cyclic hypoxia on malignant MDA-MB-231 cells. Pathway analysis revealed distinct transcriptional responses to the two hypoxic regimens. Chronic hypoxia increased epithelial-to-mesenchymal transition (EMT) signaling, and reduced inflammatory pathway activity, whereas cyclic hypoxia increased MYC target gene activation (**Figure 2A**). These findings align with our previous observations that both endogenous oscillations in HIF-1 transcriptional activity, and cyclic hypoxia permit escape from HIF-1 mediated cell-cycle arrest^12,18^. CCND1 gene, which encodes Cyclin D1, a crucial target of HIF-1 by c-Myc antagonization^7^ showed increased chromosomal accessibility and expression in cyclic hypoxia (**Figure 2B**). Similarly, Keratin-19 encoding KRT19, a direct transcriptional target of HIF-1a and a key biomarker as well as regulator of epithelial-mesenchymal transition (EMT)^34^, was expectedly increased expression in stable hypoxia, but decreased in expression even more than normoxia in cyclic hypoxia (**Figure 2B**). Transcriptional regulator analysis showed that compared with stable hypoxia, cyclic hypoxia showed intermediate activity for most regulators except for TCF7L2, ATF4, EZH2, FOXA1 (**Figure 2C**, **S3A**). These data indicate that cycling hypoxia partially reversed transcriptional programs induced by sustained hypoxia, while allowing proliferative capacity.

**Figure 2.**
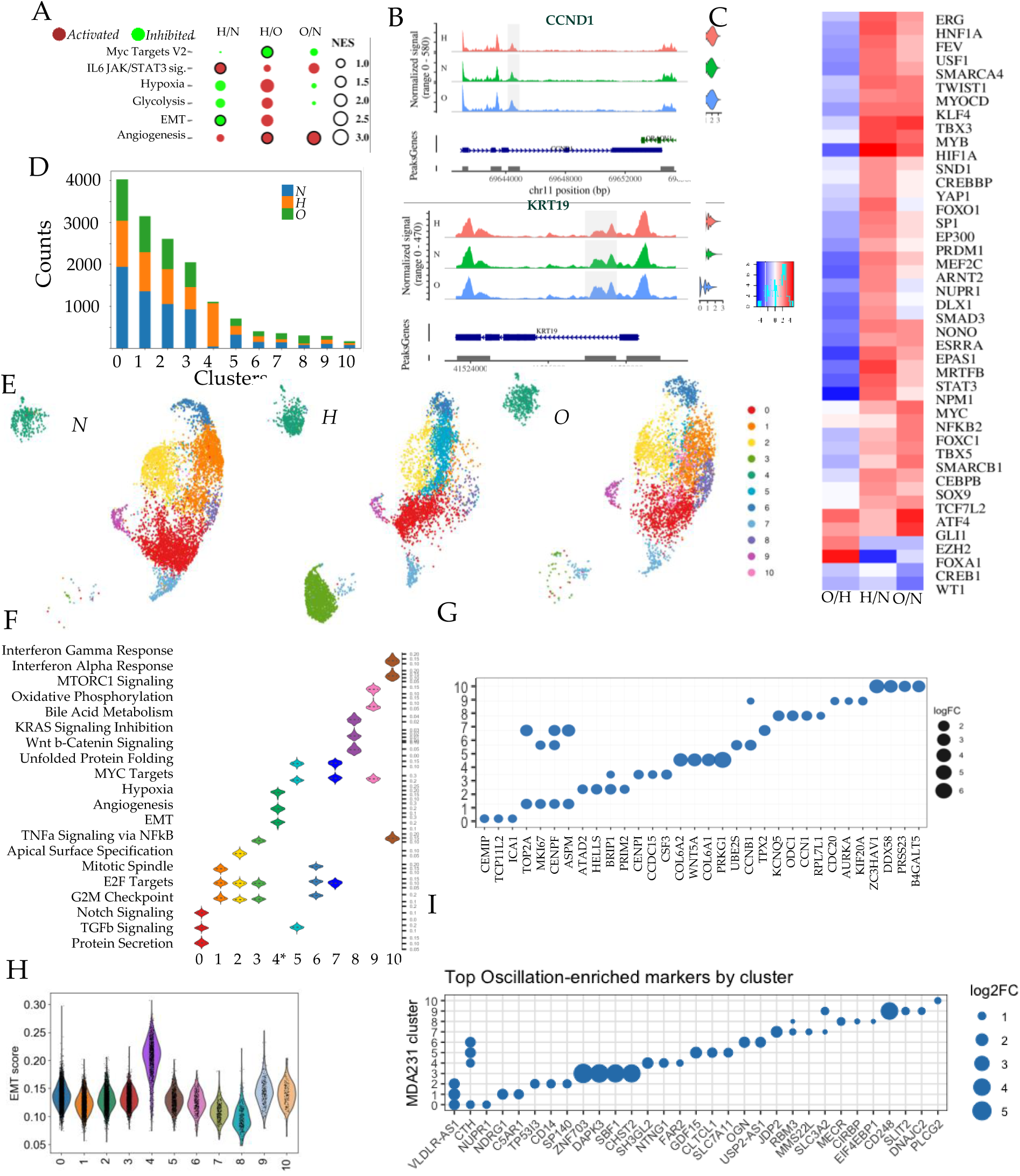
Cycling Hypoxia Results in Emergence of malignant subpopulation with high mesenchymal transcriptomic signature. (A) Relative enrichment of pathways differentially activated in MDA-MB-231 cells in normoxia (N), stable (H) and cycling hypoxia (O) relative to each other; (B) ATACseq derived peaks in the promoter regions of genes CCND1, and KRT19 in N, H, O conditions; Shown are chromosomal position (bottom), identified peaks, and violin plot of gene expression (right); (C) Upstream transcription factors based on downstream gene expression patterns differentially activated in relative conditions. (D) Relative proportions of identified subclusters in N(blue), H (orange), O (green); (E) UMAP showing subclusters identified in MDA-MB-231 cells, with the color coded legend to the right. (F) Distribution of top 3 activated pathways in each cell grouped by subclusters. (G) Top 4 marker genes for each subcluster filtered for f.c.>0.5; log(f.c.) for each gene compared in the given cluster with all other clusters; (H) EMT score for each malignant cell, shown as violin plot grouped in MDA-MB-231 subclusters; (I) Top genes differentially expressed in cyclic hypoxia (O) for each subcluster filtered for f.c.>0.5.

To determine if these transcriptional changes were accompanied by shifts in the cellular states, we observed gene expression based sub-clusters of MDA-MB-231 cells, identifying 11 distinct subpopulations (**Figure 2D**). Almost all subclusters were represented across all conditions, but three subpopulations displayed enrichment patterns specific to stable hypoxia (**Figure 2D-E**). Cluster 4 was detected nearly exclusively in stable hypoxia, while cluster 8, and to a lesser extent cluster 7 were enriched in cyclic hypoxia (**Figure 2D-E**). Pathway analysis and marker genes of these subclusters suggested distinct phenotypic characteristics for these subpopulations (**Figure 2F-G**, **S3B**). Cluster 4, which was present enriched exclusively in stable hypoxia, but was reversed in proportion in cyclic hypoxia was characterized by high expression of PRKG1, COL6A1-3 and WNT5A, and activation of EMT and angiogenesis pathways. Cluster 8, enriched in cyclic hypoxia, displayed activation of canonical WNT/β-catenin signaling, while cluster 7 was enriched for cell-cycle and unfolded protein response pathways.

To further characterize the malignant potential of cyclic hypoxia-specific subclusters, we quantified EMT state at a single cell level using a network based framework to quantify the hybrid EMT state of each cell. This score is based on the premise that cells occupy a continuum of epithelial to mesenchymal states, and regulatory feedback loops could create multistability, and consequent hybrid E/M phenotypes^35,36^. The absolute values of EMT scores for all subclusters were high, as expected for highly malignant MDA-MB-231 cells, however, cluster 4 displayed significantly higher EMT score compared to other subpopulations (**Figure 2H**). Partial EMT score also showed higher levels in cluster 4, while reduced levels in cluster 8 (**Figure S3C-D**).

Cluster 7, and 8, which were enriched in cyclic hypoxia had the lowest EMT score, which was consistent with gene expression analysis indicating high proliferative potential. Cluster 7 was characterized by TOP2A, and CENPH, classic G2/M checkpoint and mitotic markers, ASPM which is essential for mitotic spindle function orchestrating orderly cell division. In cyclic hypoxia, cluster 7 specifically enhanced expression of MMS22L crucial for DNA repair during replication, and RBM3, which is a heatshock protein known for stabilizing mRNA during high-rate proliferation (**Figure 2G,I**). Cluster 8, also enriched in cyclic hypoxia was enriched in UPR and stress pathway, and was characterized by genes critical for cell adhesion, and angiogenesis CCN1, as well as for metabolic adaptation indicated by MECR responsible for fatty acid synthesis in the mitochondria, and ODC1, a critical enzyme in polyamine synthesis (**Figure 2** **G, I**). Overall, these analyses showed that cyclic hypoxia restores stable hypoxia induced cell cycle arrest, while reversing the enrichment of a highly mesenchymal subpopulation.

### Cyclic hypoxia enhances gene expression in fibroblasts related to cell cycle and stromal inflammation

Although role of cancer associated fibroblasts as important regulators of cancer progression is well recognized, their regulatory role as being activators, or inhibitors of metastasis is still not settled^37,38^. Targeting activated CAFs for elimination has produced contradictory results in clinical trials, ushering in investigations into CAF subtypes for a more resolved understanding of stromal contribution to cancer^39,40^. We examined both stromal cell types, Fbs, and MSCs, for the transcriptional changes in sustained and cyclic hypoxia. Hypoxia significantly reduced anti-oxidant ROS pathway, and Myc target activation in Fbs indicating reduced proliferation in hypoxia (**Figure 3A**). Cycling hypoxia increased activation of UV response pathway indicating of high reactive oxygen species, reversed hypoxia induced inactivation of Myc target expression, and decreased inflammatory response signaling, opposite to the effect of sustained hypoxia (**Figure 3A**). Subpopulation examination showed that sustained hypoxia had a strong effect on bCAF gene expression, resulting in disappearance of most subclusters, and emergence of two exclusive clusters, 3 and 6 (**Figure 3B-C**). Differential gene expression patterns characterized Fb clusters, with hypoxia enriched cluster 3 showing higher expression of well known HIF-1 driven genes including ENO1, BNIP3, LDHA, and CA9 regulating glycolysis, mitophagy, lactate production from pyruvate and acidification respectively (**Figure S4**). Cycling hypoxia reversed most of the changes induced by sustained hypoxia at the cell population level, except for cluster 0, and cluster 4, which were enriched even more than in normoxia (**Figure 3B-C**). GO analysis of cluster 0 showed increased activation of collagen fibril organization, integrin signaling, and basement membrane organization, suggesting more activated CAF state (**Figure 3D**).

**Figure 3.**
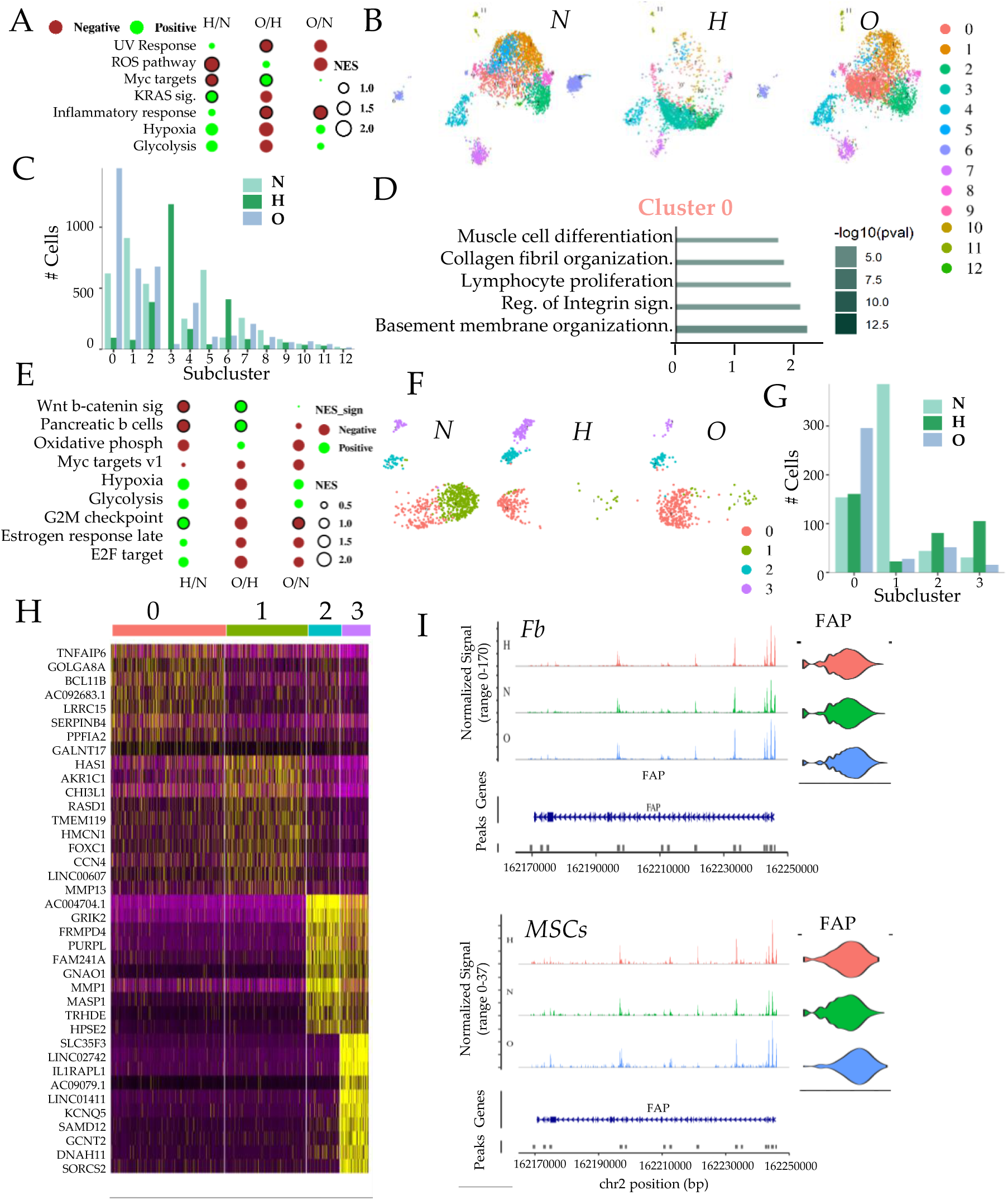
Transcriptomic responses of stromal populations in breast TME co-culture in stable and cyclic hypoxia. (A-E) Transcriptomic changes in Fibroblasts in stable and cyclic hypoxia. **(A)** Differential pathway activation in Fbs in N, H, O relative to each other; (B) t-SNE plots showing subclusters in Fbs identifying subpopulations in N, H, and O conditions, with relative proportion for each subclusters shown in (C). (D) Top pathway activation (length and direction), and p-value (thickness) for oscillation enriched Fbs cluster 4. **(E-H) Transcriptomic changes in MSCs in stable and cyclic hypoxia.** (E) Differential pathway enrichment in MSCs in N, H, O relative to each condition; (F) t-SNE plots showing MSC subclusters, with relative cell number in each condition shown in (G); (H) Top differentially expressed gene in each sub-cluster; (I) Chromosomal accessibility distribution in cis-regulatory region of FAP gene in Fbs and MSCs in N, H, O; Violin plots show distribution of FAP expression.

MSCs, another stromal cell source, home to breast cancer likely as a reparative response to the wounding of basal lamina, and inflammation caused by growing cancer^15,41^. Pathway analysis revealed that MSCs responded quite differently compared to Fbs to both sustained, and cyclic hypoxia. The effect of hypoxia on MSCs was to downregulated canonical WNT/b-catenin signaling, and activate pathways associated with cell cycle and DNA replication, including G2M checkpoint and E2F target (**Figure 3E**). Most of these changes were reversed in cycling hypoxia, which inactivated cell cycle related pathways even more than normoxia (**Figure 3E**). A resolved examination of cell subclusters showed that hypoxia resulted in depletion of cluster 1 which was unchanged in cyclic hypoxia, but also enrichment of cluster 3 (**Figure 3F**). Cyclic hypoxia, in contrast, resulted in partial reversal of changes in subpopulation by sustained hypoxia, with the exception of cluster 0, which was only specifically increased in cycling hypoxia (**Figure 3G**). Cyclic hypoxia enriched cluster 0 was characterized by higher expression of Leucine-Rich Repeat Containing 15, LRRC15, a hallmark of specific TGFβ driven myofibroblast CAF subpopulation^42,43^ (**Figure 3H**). This subpopulation has been extensively reported to be associated with ECM alignment, and therapeutic resistance^43,44^. Cluster 0 also showed increased TNFα Induced Protein 6, TNFAIP6, which encodes a secreted glycoprotein TSG-6, a critical regulator of ECM reorganization and tissue remodeling, promoting inflammatory cell infiltration ^45,46^. Other characteristic genes also indicated that cycling hypoxia enriched a subpopulation with immunomodulatory and activated fibroblast state, including BCL11B, a stroma-specific transcription factor responsible for maintenance of mesenchymal identity in stress^47^, Liprin-alpha-2 encoded by PPFIA2 which regulates focal adhesion, fibroblast spreading and matrix stiffening^48^. Together, these data suggest that cycling hypoxia activates a distinct transcriptional program in stromal cells, selectively enriching subpopulation with activated fibroblast signature. Indeed, both in Fbs and MSCs, expression and chromosomal accessibility of Fibroblast Associated Protein (FAP) was increased in cycling hypoxia (**Figure 3I**).

### Cyclic hypoxia enriches immunomodulatory macrophage subpopulation

In our multicellular model, we had added unpolarized (M0 state) macrophages, differentiated from PBMCs (**Figure S5A-B**). Although in the early stages of cancer development, tissue resident macrophages are active^25,49,50^, monocytes are recruited to the cancer, likely as a reparative response to wounding, and are quickly differentiated into macrophages^51^. Examination of subpopulation showed that sustained hypoxia relatively increased cluster 2, while cyclic hypoxia distinctly increased cluster 0 (**Figure S5C-D**). Cluster 0 was characterized by a cohesive gene expression signature associated with Tumor-Associated Macrophages (TAMs) with many genes associated with lipid metabolism, and lipid scavenging, including MSR1, LIPA, GSN, SCD, A2M, and GM2A. This characteristic gene expression indicates transformation of macrophages into “foamy” TAMs, which utilize lipid byproduct in solid cancers, and can shield the tumor from immune system^52^. In addition, higher expression of KYNU which encodes Kynureninase, was also indicative of the immunomodulatory role of cluster 0 cells, enriched in cyclic hypoxia^53^. Pathway analysis also confirmed activation of lipid peroxidation pathway, lysosomal acidification as well as tolerance induction and type II interferon pathway (**Figure S5E**).

### Cycling hypoxia elevates WNT5a/ROR1 interaction between stroma and malignant cancer cells

Our observations indicated that cycling hypoxia had a distinct influence on the cellular component in breast cancer TME, resulting in emergence of stromal and cancer subpopulations associated with mesenchymal phenotype. Cyclic hypoxia enriched subpopulation had increased Wnt signaling, so we identified if Wnt receptors were enriched in the condition. We found ROR1 expression to have specifically increased in cyclic hypoxia in single cell data (**Figure 4A**), confirmed with RT-PCR (**Figure 4B**).

**Figure 4.**
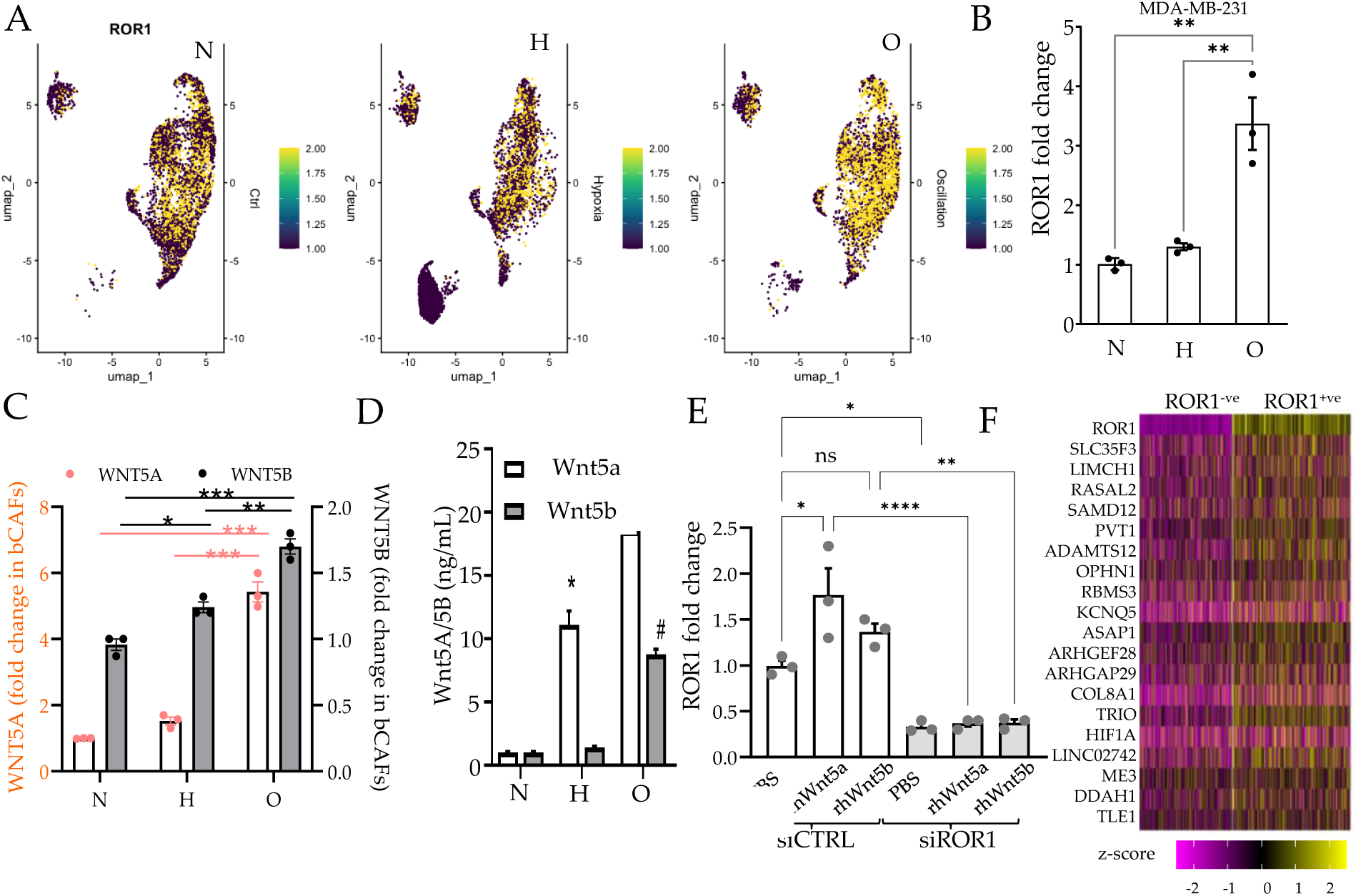
Cycling hypoxia activates Wnt5a-ROR1 axis in bCAF-TNBC paracrine interaction, driving cancer proliferation and invasion. (A) UMAP showing relative expression of ROR1 in MDA-MB-231 cells in normoxia (N), stable hypoxia (H), and cycling hypoxia (O). (B) RT-PCR showing relative expression of ROR1 in MDA-MB-231 in N, H, O; (C) RT-PCR showing relative expression of WNT5A, and WNT5B in breast cancer isolated CAFs in N, H, O conditions; (D) Secreted Wnt5a and Wnt5b measured by ELISA by bCAFs in N, H, O; (D) Relative ROR1 expression in MDA-MB-231 cells in N, H, O; (E) RT-PCR of relative ROR1 expression in MDA-MB-231 cells in the presence of recombinant Wnt5a and Wnt5b treatment, with and without prior silencing of ROR1 gene; (F) Heatmap showing top differentially expressed genes in MDA-MB-231 single cells positive and negative for ROR1 expression.

We then used CAFs isolated from breast cancer biopsies to test transcript levels of WNT5A and WNT5B, potential ligands for ROR1 receptor in MDA231. At basal levels, WNT5A and WNT5B transcripts were only present in bCAFs, while ROR1 was higher in MDA-MB-231 (**Figure S6A**). Incidentally, ROR1 is typically expressed at low levels in adult tissues, but is increased in many tumor types^54^. RT-PCR showed that both WNT5A and WNT5B transcripts increased significantly higher in stable hypoxia in bCAFs, and even further in cyclic hypoxia (**Figure 4C**), also confirmed by ELISA (**Figure 4D**).

We then tested the effect of Wnt5a and Wnt5b treatment on MDA-MB-231 cells (**Figure S6B**). Wnt5a treatment unexpectedly showed a moderate increased expression of ROR1 expression, this effect was completely diminished when ROR1 was silenced, suggesting positive feedback (**Figure S6C, 4E**). Examining differentially expressed genes in the single cell data between MDA-MB-231 cells expressing ROR1 vs those that did not identify many genes associated with higher mesenchymal state (**Figure 4F**). These included LIMCH1 and TRIO associated with cytoskeletal reorganization and invasiveness, as well as COL8A1 and ADAMTS12, suggesting remodeling of extracellular matrix. Interestingly, we also found that ROR1+ve cells expressed PVT1, which has been described to increase chemoresistance as well as proliferation^55^, as well as HIF1A levels.

### Elevated Wnt5a-ROR1 signaling between stroma-cancer in cycling hypoxia enhances tumor growth and invasiveness

What phenotypic effect could the activation of Wnt-ROR1 signaling have on cancer progression? The non-classical Wnt signaling activated by ROR1 has been implicated in many cancers, and is a primary driver of cell migration associated with poor patient prognosis^54^. Single cell differential analysis also indicated an aggressive growth and invasive phenotype in ROR1+ve cells. We therefore tested the effect of Wnt5a-ROR1 signaling on MDA-MB-231 proliferation and migration. ROR1 silencing reduced cell proliferation, secondarily measured by WST-8 metabolization in MDA-MB-231 cells (**Figure 5A**). The effect of both rhWnt5a and rhWnt5b ligands on MDA-MB-231 proliferation and their dependence on ROR1 was confirmed both by RT-PCR for MKI67 gene (**Figure 6B**), as well as nuclear localization of Ki67 (**Figure 5C-D**).

**Figure 5.**
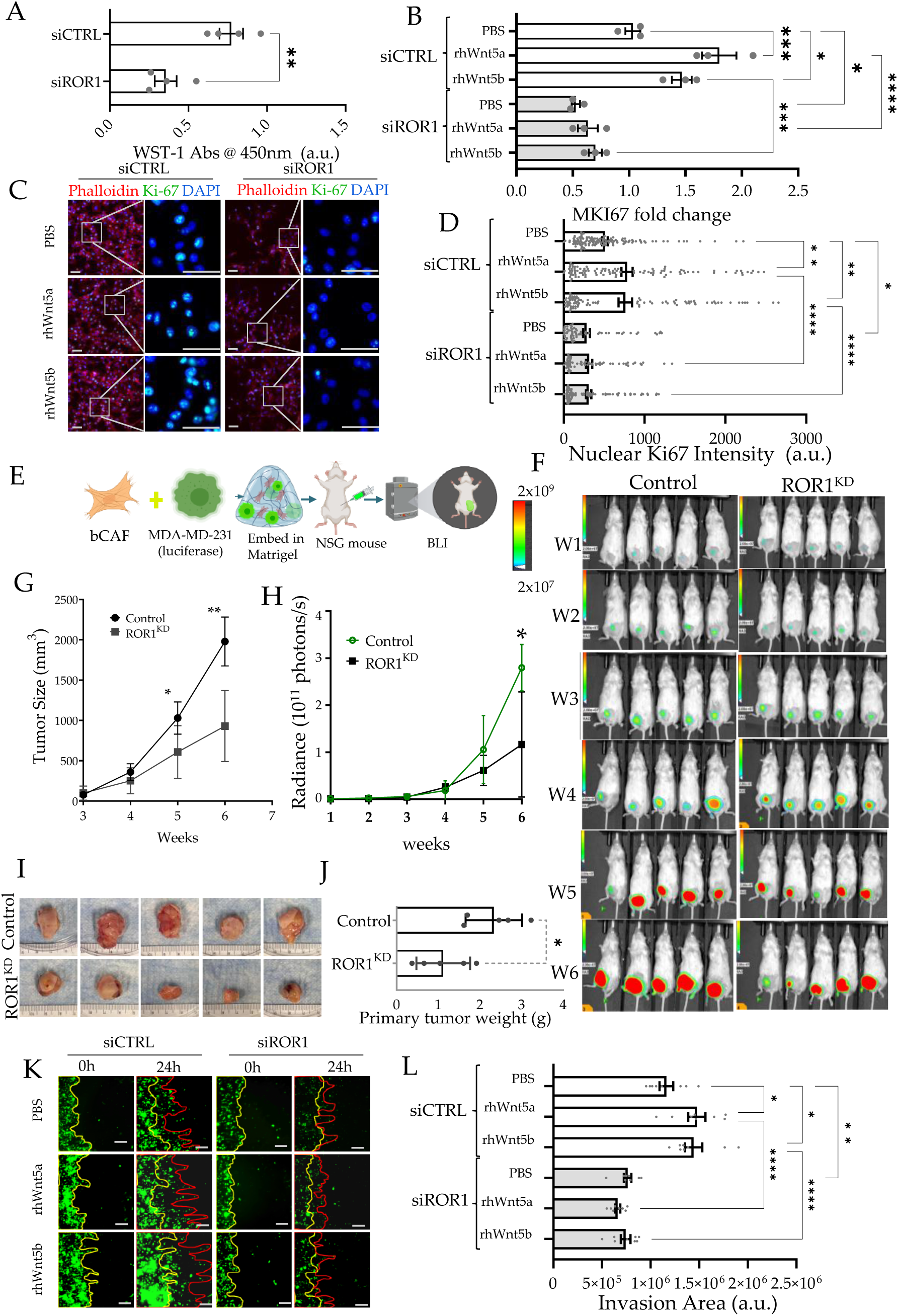
Cyclic hypoxia driven Wnt5a/b-ROR1 signaling between bCAFs and MDA-MB-231 promote tumor growth and invasion. (A) WST-1 based metabolic readout for MDA-MB-231 cells treated with recombinant human Wnt5a (rhWnt5a) 48h post ROR1 gene silencing; (B) RT-PCR showing relative MKI67 gene expression in MDA-MB-231 cells in response to ROR1 gene silencing in the presence of rhWnt5a or rhWnt5b; (C-D) Immunofluorescence based presence of nuclear Ki67 in MDA-MB-231 with or without ROR1 gene silencing, after treatment with recombinant Wnt5a, and Wnt5b, scale bar is 50 µm. (E-J) Mouse co-xenograft model of bCAFs and MDA-MB-231 silenced for ROR1; (E) Schematic showing experimental plan: Matrigel embedded orthotopic xenografting of luciferase labeled MDA-MB-231 cells with bCAFs; MDA-MB-231 cells clonally selected and silenced for vehicle (scrambled), or ROR1; (F) Bioluminescence imaging (BLI) post co-xenograft at day 1 (D1), and each subsequent week; luminescence quantified in (G); and gross tumor size measured by caliper (H); (I-J) Tumor size post resection. (K-L) Wnt5a/b driven ROR1 activation by bCAFs promotes TNBC collective invasion; (K) Representative image of collective MDA-MB-231 invasive front on underlying anisotropic nanostructured array at 0 hrs, and 24 hrs with, or without ROR1 gene silencing in the presence of recombinant Wnt5a and Wnt5b, with quantification shown in (L), scale bar is 100 µm.

We then tested the effect of ROR1 expression in MDA-MB-231 *in vivo* in an orthotopic xenograft NSG mouse model by injecting both bCAFs and luciferase expressing MDA-MB-231 cells with or without silencing ROR1 gene (**Figure S6C-D, 5E**). Bioluminescence imaging over 7 weeks showed a significantly slower tumor growth in mice engrafted with ROR1^KD^ MDA-MB-231 and bCAFs (**Figure S6B, 5F**). Both bioluminescence radiance reflecting MDA-MB-231 cell number (**Figure 6G**), and gross tumor size, reflecting malignant cancer and stroma (**Figure 5H**) grew significantly slower in the case of ROR1^KD^. Finally, because ROR1+ve cells were found to express many genes indicating enhanced mesenchymal state, we tested their invasiveness using a previously described high sensitivity stromal invasion assay^56^. Fluorescently labeled MDA-MB-231 and bCAFs were patterned orthogonally on a substrate of underlying anisotropic nanopatterned ridges, and invasion of MDA-MB-231 cells into the stromal compartment measured by live cell microscopy. We found that both Wnt5a and Wnt5b treatments increased the capacity of MDA-MB-231 to invade, but ROR1 gene silencing reduced the enhanced Wnt induced invasion (**Figure 5K-L**). Overall, these validations confirmed that cyclic hypoxia can preferentially enhance activation, and stabilization of Wnt5a-ROR1 paracrine axis between CAFs and cancer cells, promoting cancer proliferation and invasion.

## Discussion

Hypoxic niches frequently arise in tumors due to the mismatch between oxygen transport and consumption. Although, the causal factor is primarily the proliferation of cancer cells outgrowing the oxygen supply, all cellular components in the TME are exposed to the resulting oxygen deprivation. Hypoxic niches in solid tumors are not stable, and cells may experience transient, or oscillatory/cyclic hypoxia^11,57^. Cyclic hypoxia may occur even distal to blood vessels as a consequence of fluctuations in erythrocyte flux^58^. These temporal fluctuations are further compounded by endogenous oscillations in HIF-1 transcriptional activity in stable hypoxia^12,29^.

The present findings corroborate our previous observation that oscillatory/cyclic hypoxia permit cancer cells to partially evade HIF-1 induced cell cycle arrest^59^, providing a potential explanation for continued tumor growth in hypoxia despite transcriptional, and non-transcriptional inhibition of cell cycle by HIF-1^6^. Consistent with earlier studies, transcriptional response to cyclic hypoxia resembled those observed in sub-hypoxic condition^60^. Cells may integrate prolonged cyclic hypoxia as an averaged signal between hypoxic, and reoxygenation state. Such averaging of signals may account for restoration of tumor proliferation and partial reversal of chronic hypoxia induced phenotypes. However, a subset of genes responded distinctly in cyclic hypoxia, diverging from the observed response in sustained hypoxia, either more heightened response or even opposite responses. In this study, we focused mostly on these unexpected gene expression changes, which suggest a transcriptional circuitry particularly sensitive to the fluctuations in hypoxia, rather than its temporally averaged levels. The distinct gene expression patterns in cycling hypoxia could be explained by transcriptional co-regulation of downstream genes with HIF-1, and another HIF-1 regulated transcription factor, if they act in antagonistic manner, forming quasi incoherent feed forward loops^29^.

A central finding was that responses to cyclic hypoxia, differently from sustained hypoxia, were highly cell type specific. Breast CAFs displayed increased activation of mitotic pathways, and enhanced inflammatory signaling in cyclic hypoxia, in contrast to the suppressive effect in chronic hypoxia. MSCs exhibited a particularly pronounced response to cyclic hypoxia. Whereas, Wnt/b-catenin signaling was suppressed in hypoxia, it was restored in cyclic hypoxia. MSCs are typically recruited to cancer in response to the wounding and tissue damage signals emanating from tumor proper to initiate regenerative programs under hypoxia. Consistent with the biology of MSCs, hypoxia led to activation of proliferative pathways in MSCs, in contrast to other cell types. In contrast, cyclic hypoxia inhibited cell cycle in MSCs. Similarly, cyclic hypoxia appeared to partially reverse sustained hypoxia induced immunomodulatory phenotype in macrophages.

We found that cyclic hypoxia appears to coordinate WNT5A and WNT5B production in stromal populations and receptor ROR1 expression in malignant cells, potentially amplifying the WNT5A-ROR1 signaling in the tumor ecosystem. Wnt signaling is critical for mammary gland development, and aberrant Wnt signaling is common, and rather a characteristic of TNBC^61^. Wnt5a, as well as Wnt5b increased MDA-MB-231 proliferation in ROR1 dependent manner, both *in vitro*, and also in a human cancer-CAF co-xenograft model. We also demonstrated that WNT5A-ROR1 signaling axis increases the invasive capability of malignant cells to invade the stromal compartment. ROR1 expression is absent in normal breast cell, and its increase in TNBC is associated with poor survival^62^. ROR1 levels increased in the presence of Wnt5a and Wnt5b, suggesting that maintenance of higher ROR1 levels in the tumor ecosystem could also be strengthened by this small subpopulation.

Overall, our findings suggest that cyclic hypoxia is not merely an intermediate between normoxia and hypoxia, but is decoded differently within the TNBC cellular microenvironment with specific consequences, including in readjustments of the cellular state composition, and coordinated reprogramming of pro-tumorigenic WNT5A-ROR1 signaling.

## Authors’ contribution

Y.L. conducted most experiments, developed methods and performed data analysis,

S.L. performed most bioinformatics analysis;

S.S. and M.K.J. performed part bioinformatics analysis,

X.W. assisted in animal experiments,

W.D. performed invasion experiments,

D.E.E. provided part infrastructure support,

J.A. and Y.S. supervised preparation of scRNAseq and bioinformatics analysis respectively, K. conceptualized, supervised and drafted the manuscript.

## Ethics Statement

All animal studies were performed under an protocol approved by the Institutional Animal Care and Use Committee (IACUC) at UConn Health. Protocol Number AP-200781 approved December 06, 2024.

## Acknowledgements

We gratefully acknowledge the support of our funding agencies: National Cancer Institute (1R37CA248161), and American Heart Association predoctoral fellowship program (24PRE1196464), Param Hansa Philanthropies. We also acknowledge the Single Cell Biology service and Cyberinfrastructure high performance computing resources at The Jackson Laboratory supported in part by the JAX Cancer Center (P30 CA034196).

## Data Availability Statement

All data are made available to reviewers or readers on request. scRNAseq and ATACseq data are available on figshare: https://doi.org/10.6084/m9.figshare.28408919

**Acknowledgements:**

