## Supplementary Information for "Cyclic Hypoxia drives Wnt5a-ROR1 stroma-tumor crosstalk promoting breast cancer growth"

**Supplementary Figures**

**Supplementary Figure 1. Differential gene expression in stable and cyclic hypoxia in cell types.** (A) Post-analysis counts of each cell type in Normoxia (N), Hypoxia (H), and cycling Hypoxia (O). Heatmaps showing z-score of top differential genes in normoxia (N), stable hypoxia (H), and cyclic hypoxia (O) in each cell for (B) MDA-MB-231, (C) bCAFs, (D) MSCs, (E) Macrophages, and (F) HUVECs in co-culture.

**Supplementary Figure 2. Differential activation of Transcription Factors in stable, and cyclic hypoxia in cell types.** Scatterplots showing IPA based activation score of upstream transcriptional regulators in stable hypoxia vs normoxia (x-axis), and cyclic hypoxia vs stable hypoxia (y-axis) for (A) MDA-MB-231, (B) bCAFs, (C) MSCs, (D) Macrophages, and (E) HUVECs. TFs with significant activation scores (<0.05) are labeled in blue.

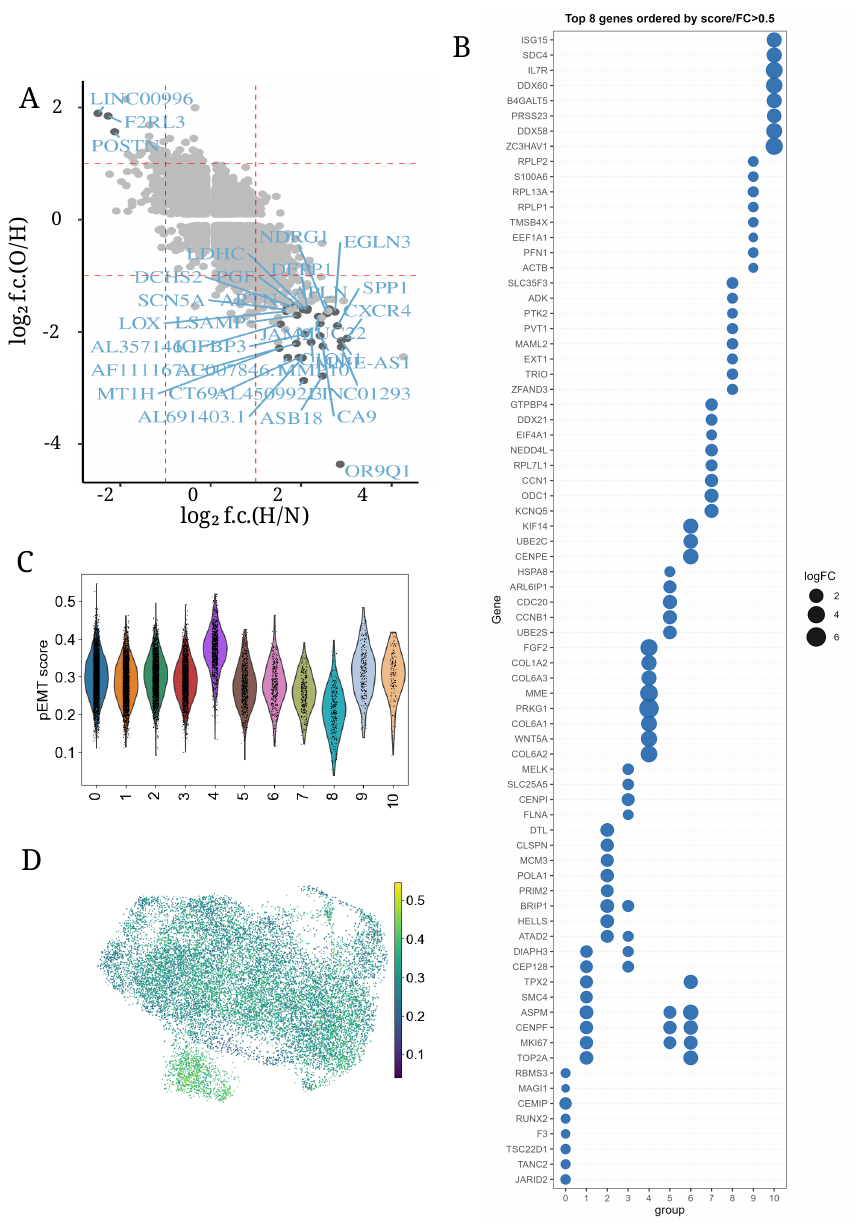

**Supplementary Figure 3. Subcluster analysis of MDA-MB-231 cells. (A) Scatter plot showing** differential expression of transcription factor encoding genes shown as log2(fold-change) between stable hypoxia (H) and normoxia (N) in x-axis, and cyclic hypoxia (O) vs stable hypoxia (H) in y-axis; TFs with significantly high fold-change are labeled. (B) Bubble plot showing log2(fold-change) of top 8 genes in each sub-cluster compared to mean expression in all other sub-clusters; (C) Violin plot showing partial EMT score calculated for each single MDA-MB-231 cell; (D) UMAP of MDA-MB-231 cells showing partial EMT score (pEMT) of individual cells.

**Supplementary Figure 4. Differential gene expression in bCAF cells.** Heatmap showing expression of top differential genes in each identified sub-cluster.

**Supplementary Figure 5. scRNAseq analysis of stable and cyclic hypoxia on macrophages in TME co-culture (A-B) Characterization of PBMCs derived macrophages. (A)** Phase-contrast image showing macrophages 7 days after differentiation from monocytes; (B) Flow cytometry showing CD11c and CD14 expression in differentiated, unpolarized macrophages. **(C)** t-SNE plots showing clusters of initially unpolarized macrophages in co-culture with other cell types in TNBC TME in normoxia (N), stable hypoxia (H), and cyclic hypoxia (O), with relative proportion in each condition shown in (D). (E) Heatmap showing expression of top differential genes in sub-clusters; (F) Selected pathways in cyclic hypoxia enriched sub-cluster 0.

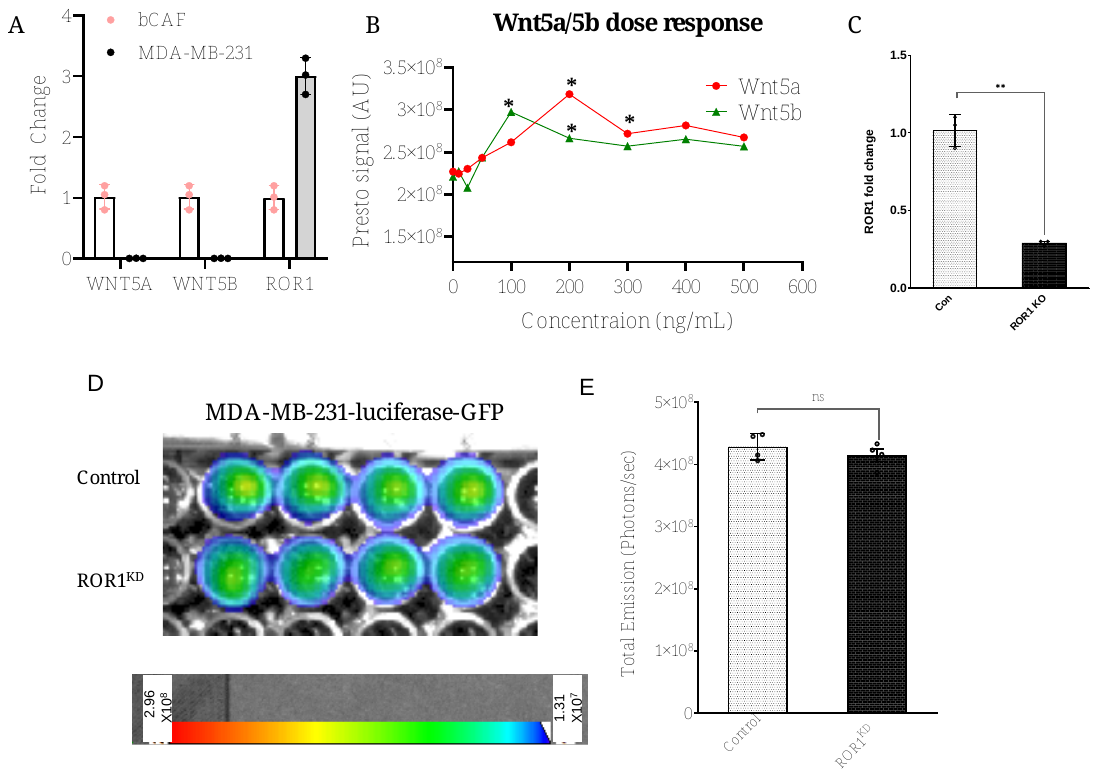

**Supplementary Figure 6. Wnt5a and Wnt5b experimental characterizations.** (A) Relative expression of WNT5A, WNT5B, and ROR1 transcripts by RT-PCR in bCAFs and MDA-MB-231; **(B)** Dose response of Wnt5a and Wnt5b treatment on MDA-MB-231 to identify working range concentration for an effect on cell viability and proliferation. (D) Confirmation of ROR1 silencing in clonally selected cells by RT-PCR. (D-E) Confirmation of no biased change in luciferase signal in ROR1KD vs control (scrambled silenced) cells.
